# Evidence of massive leukemic progenitor expansion in blood during molecular recurrence after TKI discontinuation in chronic myeloid leukemia (CML)

**DOI:** 10.1101/805648

**Authors:** Ali G Turhan, Patricia Hugues, Nathalie Sorel, Christophe Desterke, Jean-Henri Bourhis, Annelise Bennaceur-Griscelli, Jean-Claude Chomel

## Abstract

Chronic myeloid leukemia (CML) represents one of the major success stories of targeted therapies of the 21st century with the use of tyrosine kinase inhibitors (TKIs). Following discontinuation of TKIs in deep molecular remission context, at least half of the patients experience molecular relapse. Cellular events occurring during this phase are not yet established. In this work, we show a massive amplification of clonogenic progenitors or CFCs (colony-forming cells) in the peripheral blood of patients in molecular recurrence after a TKI discontinuation. We demonstrate, by qRT-PCR analysis on individual and pooled CFCs, that leukemic clonogenic progenitor expansion in the peripheral blood represents the first cellular event before any evidence of cytological change. These findings also suggest that the amplification of the leukemic clone during the first stages of CML takes place not only in the bone marrow but also in the peripheral blood.

## INTRODUCTION

Tyrosine kinase inhibitor (TKI) cessation strategies have shown both their great interest and their feasibility for selected patients with chronic myeloid leukemia (CML) [1]. It has been clearly demonstrated that TKI therapy (Imatinib, Nilotinib, Dasatinib) can be safely stopped for patients in sustained deep molecular response or DMR (at least one year in molecular response > MR^4^). However, in such a situation, 50-60% of patients experience molecular recurrence [2, 3]. The loss of major molecular response or MMR (*BCR-ABL1/ABL1*^IS^ > 0.1%) is now considered as the main criterion for evidence of molecular relapse and restarting TKI therapy [4]. In the majority of cases, molecular recurrence occurs within the first year of therapy cessation (usually in the first six months). The loss of MMR in such a short time is compatible with the resurgence of leukemic hematopoiesis from residual leukemic stem cells (LSCs) or progenitors. Indeed, we and others have previously demonstrated the persistence of *BCR-ABL1*-positive progenitors or stem cells in bone marrow from CML patients in DMR [5, 6]. Using CD26 expression, it has recently been shown that LSCs can persist in the peripheral blood of CML patients in molecular response, including those in treatment-free remission [7]. However, the presence of residual leukemic progenitors or stem cells, detected by CFC (colony-forming cell) or LTC-IC (Human Long-Term Culture-Initiating Cell) assays, has not been consistently associated with a molecular relapse after Imatinib cessation [8]. Although long-term persistence of a leukemic progenitor/stem cell reservoir is likely to represent the primary cause of molecular recurrence after TKI discontinuation, cellular dynamics of these early relapses and the pattern of cells giving rise to them have yet to be determined. For that purpose and based on a few selected patients, we wondered whether leukemic progenitor cells can be detected in the peripheral blood at the time of molecular recurrence, and what could be their impact in this critical event.

## METHODS

### In vitro hematopoietic progenitor and stem cell assays

Mononuclear cells (MNCs) were isolated from blood and bone marrow samples on Histopaque density gradient separation (Sigma-Aldrich, St Louis, MO). CFC assays were performed by plating 50,000 MNCs on semisolid methylcellulose Methocult H4435 medium (StemCell Technologies, Vancouver, Canada). At day +14 of culture, CFU-GM (colony forming unit-granulocyte macrophage), BFU-E (burst forming unit-erythroid) and CFU-GEMM (colony forming unit-granulocyte erythrocyte monocyte macrophage) were enumerated, plucked from methylcellulose and put (individually or by pools of 2-5 colonies) into RNA extraction buffer (Arcturus Bioscience Inc, Mountain View, CA).

### RNA extraction and cDNA synthesis

Total RNA was extracted from individual and pooled colonies using the PicoPure RNA isolation kit (Arcturus Bioscience Inc) according to the manufacturer’s instructions. RNA was reverse transcribed using the High Capacity cDNA Reverse Transcription Kit (Applied Biosystems, Foster City, CA).

### Detection of *BCR-ABL1* mRNA on individual and pooled colonies

The presence of *BCR-ABL1* mRNA transcripts was analyzed by qRT-PCR according to ELN recommendations. Each plate consisted of cDNA samples from hematopoietic colonies, positive and negative controls and each assay was done in duplicate. *ABL1* amplification was used to assess the presence of amplifiable cDNA. In the latter case, the *BCR-ABL1/ABL1* ratio was determined using FusionQuant standards (Qiagen, Marseille, France) according to the Europe Against Cancer protocol. All results were expressed as the *BCR-ABL1/ABL1*^IS^ ratio, normalized to the International Scale by a specific conversion factor (based on a primary international reference material).

## RESULTS

We had the opportunity to study three patients (P1-P3) who presented molecular recurrence following Imatinib discontinuation after obtaining a deep molecular response (MR^4.5^) for at least two years. All patients were diagnosed in the chronic phase and were treated with Imatinib 400 mg/day as first-line therapy. Durations of TKI therapy and DMR before discontinuation were 2.5-15 and 2-6 years respectively (Table 1). For patient P1, a CFC assay was performed in a bone marrow aspirate before Imatinib discontinuation showing no *BCR-ABL1*-expressing progenitors (the methodology is briefly described below). At the time of molecular recurrence, all patients had a normal clinical examination with normal complete blood counts and no evidence of circulating immature blood cells. Molecular relapses occurred at 2 months (*BCR-ABL1/ABL1*^IS^ = 27%), 3 months (*BCR-ABL1/ABL1*^IS^ = 13.1%), and 9 months (*BCR-ABL1/ABL1*^IS^ = 2.7%.) for patients P1, P2 and P3 respectively. Another patient (P4) with secondary resistance to second-line Nilotinib was also studied as a reference. He received TKI (Imatinib, then Nilotinib) for six years and achieved MR^4.5^ for three years before the occurrence of a sudden and significant increase in the *BCR-ABL1* ratio (10%).

CFC assays were performed from the same blood samples in which the molecular recurrence (patients P1-P3) or secondary resistance (patient P4) was documented. At day +14, clonogenic cells (CFU-GM, BFU-E, CFU-Mix) were plucked from methylcellulose, and *BCR-ABL1* mRNA expression was determined by qRT-PCR as previously reported [5]. In total, 203 CFCs were analyzed individually or in pools of 2-5 colonies. For plated 5×10^4^ peripheral blood mononuclear cells (PBMCs), the mean number of CFCs generated per dish varied from 12 to 29. The presence of *BCR-ABL1*-expressing CFCs in peripheral blood was demonstrated in all four patients (Table 1). In patient P1 and P2, all CFCs were positive (mean *BCR-ABL1/ABL1*^IS^ 1.2 and 15.9, respectively), while only four CFC pools (out of twenty) expressed *BCR-ABL1* in patient P3 (mean *BCR-ABL1/ABL1*^IS^ 0.08%). Finally, only one CFC pool appeared positive in patient P4 (mean *BCR-ABL1/ABL1*^IS^ 1.5%). When a pool is positive, the number of *BCR-ABL1*-expressing CFCs cannot be clearly specified, but only estimated. Keeping in mind the estimated number of CFCs per ml and the number of positive pooled CFCs, the theoretical number of leukemic CFCs at the time of molecular recurrence was 467±139/ml, 374±93/ml of peripheral blood for patients P1 and P2 presenting early molecular recurrence. Patient P3 who experienced a late relapse had 59±31 leukemic circulating CFCs per ml. A lower level (19±12/ml) was observed in patient P4 (on-therapy) at the time of resistance to Nilotinib. All patients had a favorable response after resuming Imatinib (patients P1-P3) or switching to Bosutinib (patient P4).

## DISCUSSION

Since a large number of CFCs are required for analysis, this type of study cannot be based on large cohorts of patients so as to highlight statistical trends. However, using a limited number of selected patients, some relevant data have appeared. As previously reported, *BCR-ABL1/ABL1* ratios appear low and heterogeneous in CFCs [9, 10]. Amongst the patients in Imatinib cessation, two presented an early molecular relapse (2-3 months) while the third had a later relapse (9 months). All of them displayed a substantial number of *BCR-ABL1/ABL1*-expressing CFCs in their peripheral blood at the time of recurrence. Based on our data, up to 2-3×10^6^ (patients with early molecular recurrence) and 4×10^5^ (patient with late molecular recurrence) CML progenitors were present in the whole peripheral blood compartment at the time of molecular recurrence. It should be noted that the large number of CFCs was detected in the absence of any abnormalities in the blood cell count.

It has been shown that Imatinib can induce the migration of CML stem cells into the bone marrow niche, thereby inducing their quiescence [11]. Furthermore, the BCR-ABL oncoprotein is known to promote cell cycle entry of LSCs that display high proliferation rates in both peripheral blood and bone marrow [12, 13]. The loss of their quiescence status could then promote their escape from the bone marrow microenvironment and initiate leukemic expansion. In theory, Imatinib cessation could lead to the release of quiescent residual LSCs from the niche and the mobilization of cycling LSCs. Interestingly, in one of our patients (P1), CFC assays performed from bone marrow four months before Imatinib discontinuation showed no evidence of *BCR-ABL1*-expressing CFCs (Table 1). This result suggests that rapid and massive expansion of a quiescent LSC clone generating leukemic CFCs migrating to peripheral blood occurred at the time of molecular recurrence.

Consequently, it may be assumed that leukemic circulating CFCs represent the first abnormal cellular compartment detectable during molecular recurrence after TKI cessation without evidence of hematological relapse. This fact also suggests that clonogenic relapse in peripheral blood precedes myeloid expansion. Based on these results, a hypothetical model explaining early molecular recurrences following TKI discontinuation can be designed (Fig. 1). Imatinib discontinuation can, in some patients, lead to the activation of LSCs (a) generating the amplification of CFCs in the bone marrow (b), which are massively mobilized into the blood where they expand (c). In this scheme, clonogenic relapse coincides with the detection of molecular recurrence by qRT-PCR (d) and precedes an eventual hematological relapse related to the proliferation of more mature leukemic cells (e-f). Of course, monthly molecular monitoring of patients in TKI cessation allows, upon loss of MMR, the resumption of targeted therapy, thereby preventing hematological relapse (Fig. 1). It remains to be determined whether some LSCs participate in this process to egress from the bone marrow to peripheral blood generating circulating CFCs (c’). To address this issue, appropriate tests such as LTC-IC assays would be needed. However, this method would require a large volume of peripheral blood in order to eventually detect a small number of LSCs. Therefore, it does not seem applicable in such a case.

**Fig. 1:**
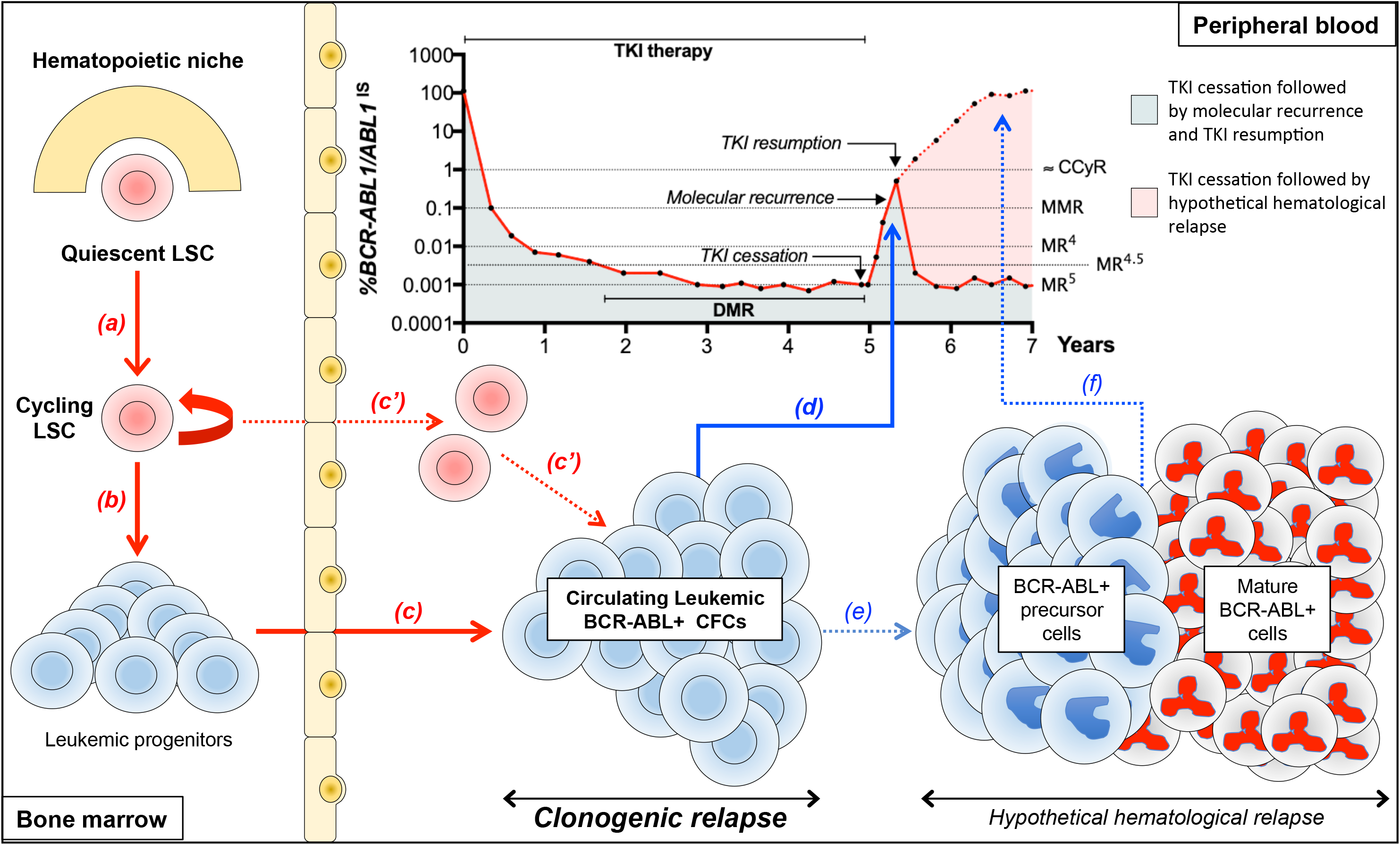
Schematic representation of the sequence of cellular events during molecular recurrence following TKI cessation. This model describes a potential sequence of cellular events leading to molecular recurrence after TKI cessation. All data are represented on a theoretical *BCR-ABL1/ABL1* molecular follow-up. (a) activation of residual quiescent LSCs upon TKI cessation. (b) generation of leukemic progenitors in bone marrow. (c) migration and massive amplification of leukemic CFCs in peripheral blood. (c’) hypothetical participation of LSCs directly migrating from bone marrow to peripheral blood to the CFC amplification process. (d) clonogenic recurrence precedes hematological relapse prevented by resuming TKI therapy. (e) potential progression of the leukemic disease (leukemic precursors and mature cells) in the absence of TKI resumption leading to a hematological relapse (f). LSC, leukemic stem cell. TKI, tyrosine kinase inhibitor; CFC, colony-forming cell; CCyR, complete cytogenetic response; MMR, major molecular response; DMR, deep molecular response (MR^4^, MR^4.5^, MR^5^).

Overall, the data reported here provide a unique opportunity to revisit the first stages of the disease which are obviously not detectable in a patient with “smoldering” CML whose diagnosis will be made only after amplification of the neoplastic clone. The first steps of CML could consequently involve leukemic progenitor egress from bone marrow to peripheral blood, and then progenitor expansion not only in bone marrow but also in peripheral blood, giving rise to an overt myeloproliferative neoplasm. Besides shedding light into the pathophysiology of CML, our findings could be of significant interest to design novel biological follow-up protocols after TKI discontinuation.

## Supporting information

Turhan et al Table 1

## CONFLICT OF INTEREST

The authors declare no conflict of interest.

## ACKNOWLEDGEMENTS

The English of the manuscript was reviewed by Jeffrey Arsham, an American medical translator.

## AUTHOR’S CONTRIBUTIONS

A.G.T. designed the study, provided patient blood and bone marrow samples, analyzed the data and contributed to manuscript writing. P.H. performed experiments. N.S. performed experiments and revised the manuscript. C.D. analyzed the data. J.H.B. provided CML patient blood samples and revised the manuscript. A.B.G. designed the study and revised the manuscript. J.C.C. analyzed the data and contributed to manuscript writing. All authors read and approved the manuscript in its final version.

## FUNDING

This research received no external funding.

## REFERENCES

1. Cortes J, Rea D, Lipton JH. Treatment-free remission with first- and second-generation tyrosine kinase inhibitors. Am J Hematol. 2019;94:346–57.

2. Etienne G, Guilhot J, Rea D, Rigal-Huguet F, Nicolini F, Charbonnier A, et al. Long-Term Follow-Up of the French Stop Imatinib (STIM1) Study in Patients With Chronic Myeloid Leukemia. J Clin Oncol. 2017;35:298–305.

3. Saussele S, Richter J, Guilhot J, Gruber FX, Hjorth-Hansen H, Almeida A, et al. Discontinuation of tyrosine kinase inhibitor therapy in chronic myeloid leukaemia (EURO-SKI): a prespecified interim analysis of a prospective, multicentre, non-randomised, trial. Lancet Oncol. 2018;19:747–57.

4. Rousselot P, Charbonnier A, Cony-Makhoul P, Agape P, Nicolini FE, Varet B, et al. Loss of major molecular response as a trigger for restarting tyrosine kinase inhibitor therapy in patients with chronic-phase chronic myelogenous leukemia who have stopped imatinib after durable undetectable disease. J Clin Oncol. 2014;32:424–30.

5. Chomel JC, Bonnet ML, Sorel N, Bertrand A, Meunier MC, Fichelson S, et al. Leukemic stem cell persistence in chronic myeloid leukemia patients with sustained undetectable molecular residual disease. Blood. 2011;118:3657–60.

6. Chu S, McDonald T, Lin A, Chakraborty S, Huang Q, Snyder DS, et al. Persistence of leukemia stem cells in chronic myelogenous leukemia patients in prolonged remission with imatinib treatment. Blood. 2011;118:5565–72.

7. Bocchia M, Sicuranza A, Abruzzese E, Iurlo A, Sirianni S, Gozzini A, et al. Residual Peripheral Blood CD26^+^ Leukemic Stem Cells in Chronic Myeloid Leukemia Patients During TKI Therapy and During Treatment-Free Remission. Front Oncol. 2018;8:194.

8. Chomel JC, Bonnet ML, Sorel N, Sloma I, Bennaceur-Griscelli A, Rea D, et al. Leukemic stem cell persistence in chronic myeloid leukemia patients in deep molecular response induced by tyrosine kinase inhibitors and the impact of therapy discontinuation. Oncotarget. 2016;7:35293–301.

9. Kumari A, Brendel C, Hochhaus A, Neubauer A, Burchert A. Low BCR-ABL expression levels in hematopoietic precursor cells enable persistence of chronic myeloid leukemia under imatinib. Blood. 2012;119:530–39.

10. Chomel JC, Sorel N, Guilhot J, Guilhot F, Turhan AG. BCR-ABL expression in leukemic progenitors and primitive stem cells of patients with chronic myeloid leukemia. Blood. 2012;119:2964–65.

11. Jin L, Tabe Y, Konoplev S, Xu Y, Leysath CE, Lu H, et al. CXCR4 up-regulation by imatinib induces chronic myelogenous leukemia (CML) cell migration to bone marrow stroma and promotes survival of quiescent CML cells. Mol Cancer Ther. 2008;7:48–58.

12. Krämer A, Löffler H, Bergmann J, Hochhaus A, Hehlmann R. Proliferating status of peripheral blood progenitor cells from patients with BCR/ABL-positive chronic myelogenous leukemia. Leukemia. 2001;15:62–68.

13. Bruns I, Czibere A, Fischer JC, Roels F, Cadeddu RP, Buest S, et al. The hematopoietic stem cell in chronic phase CML is characterized by a transcriptional profile resembling normal myeloid progenitor cells and reflecting loss of quiescence. Leukemia. 2009;23:892–99.

