## Supplementary material for "Evidence of massive leukemic progenitor expansion in blood during molecular recurrence after TKI discontinuation in chronic myeloid leukemia (CML)": Turhan et al Table 1

**Table 1**: **Extensive analysis of CFC assays in patients in molecular recurrence following Imatinib discontinuation.**

TKI, tyrosine kinase inhibitor; DMR, deep molecular response; BMC, bone marrow mononuclear cell; CFC, colony-forming cell; PBMC, peripheral blood mononuclear cell; ND, not done; NA, not applicable. * The range was estimated using the estimated number of CFCs per ml of peripheral blood and the number of *BCR-ABL1*-expressing CFCs analyzed individually or in pools.

|  | **Patients** |  | **P1** | **P2** | **P3** |  | **P4** |
| --- | --- | --- | --- | --- | --- | --- | --- |
| **Patients characteristics** | **Age /sex** |  | 47 / M | 66 / F | 87 / M |  | 83 / M |
|  | **TKI therapy** |  | Imatinib  400 mg/day | Imatinib  400 mg/day | Imatinib  400 mg/day |  | Imatinib  400 mg/day |
|  |  |  |  |  |  |  | Nilotinib  300 mg x 2/day |
| **DMR achievement (>MR^4.5^)** | |  | *ON-THERAPY* | | | | |
| **Duration** | **TKI therapy (years)** |  | 14 | 15 | 2.5 |  | 6 |
|  | **DMR duration (years)** |  | 6 | 5 | 2 |  | 3 |
| **CFC assays before cessation**  ***(in bone marrow)*** | **Volume sample (ml)** |  | 3.5 | *ND* | *ND* |  | *NA* |
|  | **BMC number (10^6^)** |  | 30 | *ND* | *ND* |  | *NA* |
|  | **CFC pools analyzed** |  | 20 CFCs  *18 individual CFCs*  *1 pool of 2 CFCs* | *ND* | *ND* |  | *NA* |
|  | **CFC pools expressing *BCR-ABL1*** |  | 0 | *ND* | *ND* |  | *NA* |
| **Treatment cessation** | |  | *OFF-THERAPY* | | |  | *ON-THERAPY* |
| **Molecular recurrence** | **Delay**  **(months)** |  | 2 | 3 | 9 |  | *NA* |
|  | ***BCR-ABL1/ABL1*^IS^ in peripheral blood** |  | 27% | 13.1% | 2.7% |  | 10% |
| **CFC assays at molecular recurrence**  ***(in peripheral blood)*** | **Volume sample (ml)** |  | 13.5 | 10 | 16 |  | 16 |
|  | **PBMC number (10^6^)** |  | 20 | 18 | 14 |  | 30 |
|  | **CFCs generated per dish for 5x10^4^ PBMCs plated** |  | 20.5 | 13 | 29 |  | 12 |
|  | **Estimated number of CFCs per ml** |  | 607 | 468 | 507.5 |  | 450 |
|  | **CFC analyzed individually or in pools** |  | 37 CFCs  *5 individual CFCs*  *13 pools of 2 CFCs*  *2 pool of 3 CFCs* | 35 CFCs  *10 individual CFCs*  *10 pools of 2 CFCs*  *1 pool of 5 CFCs* | 73 CFCs  *9 pools of 3 CFCs*  *9 pools of 4 CFCs*  *2 pool of 5 CFCs* |  | 58 CFCs  *2 individual CFCs*  *2 pools of 2 CFCs*  *6 pools of 3 CFCs*  *6 pools of 4 CFCs*  *2 pool of 5 CFCs* |
|  | **Individually or pooled CFCs expressing**  ***BCR-ABL1*** |  | *5 individual CFCs*  *13 pools of 2 CFCs*  *2 pool of 3 CFCs* | *10 individual CFCs*  *10 pools of 2 CFCs*  *1 pool of 5 CFCs* | *3 pools of 3 CFCs*  *1 pool of 4 CFCs* |  | *1 pool of 4 CFCs* |
|  | **Median *BCR-ABL1/ABL1*^IS^** |  | 1.2% | 15.9% | 0.08% |  | 1.5% |
|  | **Estimated number of leukemic CFCs per ml*** |  | 328-607 | 281-468 | 28-90 |  | 7-31 |
| **TKI resumption** | |  | *ON-THERAPY* | | | | |
| **Retreatment** | **TKI therapy** |  | Imatinib  400 mg/day | Imatinib  400 mg/day | Imatinib  400 mg/day |  | Bosutinib  300 mg/day |
|  | **Evolution** |  | MR^4^  at 18 months | MR^4^  at 24 months | MR^4^  at 12 months |  | MR^4^  at 12 months |
